# Large-scale prediction of key dynamic interacting proteins in multiple cancers

**DOI:** 10.1101/2020.12.04.411173

**Authors:** Jifeng Zhang, Xiao Wang, Zhicheng Ji, Weidong Tian

## Abstract

Tracking cancer dynamic protein-protein interactions(PPIs) and deciphering their pathogenesis remain a challenge. Here, we presented a dynamic PPIs’ hypothesis: permanent and transient interactions might achieve dynamic switchings from normal cells to malignancy, which could cause maintenance functions to be interrupted and transient functions to be sustained. Based on the hypothesis, we first predicted more than 1,400 key cancer genes (KCG) by applying PPI-express we proposed to 18 cancer gene expression datasets. Two prominent functional characteristics, “Cell cycle-related” and “Immune-related”, were presented, suggesting that it might be a general characteristic of KCG. We then further screened out key dynamic interactions (KDI) of cancer based on KCG and transient and permanent interactions under both conditions. We found that, compared to permanent to transient KDI pairs (P2T) in the network, transient to permanent (T2P) have significantly higher edge betweenness (EB), and P2T pairs tending to locate intra-functional modules may play roles in maintaining normal biological functions, while T2P KDI pairs tending to locate inter-modules may play roles in biological signal transduction. It was consistent with our hypothesis. Also, we analyzed network characteristics of KDI pairs and their functions. Our findings of KDI may serve to understand and explain a few hallmarks of cancer.

## Introduction

The dynamic interactions in cancer proteins that are reproduced systematically remain one of the challenges of cancer research. However, using genome-scale and diverse data sources, bioinformatics may be promising and impartial tool for studying the dynamic interactions of cancer proteins [1]. Protein interactions are often not “static” and interact with each other to complete a variety of biological processes. For example, proteins perform the extracellular signal transmission to achieve the external environment response through the dynamic signal network [2]. In normal cells, there is a difference in the interaction time between proteins. According to the time of action (or the lifetime of complex), the interactions between proteins can be divided into permanent interactions and transient interactions[3]. Enzyme-inhibitor, antibody-antigen, and receptor-ligand interactions in PPIs possess a high affinity, and the complexes are usually permanent (e.g., enzyme-inhibitor: thrombin-rodniin)[4], other proteins activate downstream proteins to form complexes, which are then separated, to achieve signal transduction and these complexes are considered to be transient interactions (e.g., cell proliferation complex: Ras–RasGAP)[5,6]. Dynamic protein interactions can reflect temporal and spatial changes in protein action[7,8]. In some special cases, the dynamic changes may manifest as a dramatic switching between transient and permanent PPI. And similar features of complex interface may provide a structural basis for the switching [3].

Interacting pairs in HPRD [9] or STRING [10] can only represent the interaction of proteins in a state. Exploring the dynamic interactions of proteins may help us to better understand human cancer [11]. However, only few studies of dynamic interactions of cancer proteins have been reported. For instance, Taylor et al. [12] found that dynamic modularity of the human interactome could be used to predict breast cancer prognosis through the analysis of network hub proteins. Similarly, another study analyzed glioma outcome by finding dynamic modularity in gene co-expression networks [10]. Liu et al. [13] divided protein interactions into cooperation, competition, redundancy, and dependency based on their gene expression levels, and predicted dysregulated interactions and pathways of breast cancer and pancreatic cancer using GIENA (gene interaction enrichment and network analysis). The lack of real-time structural monitoring data limits the large-scale prediction of protein interactions[14]. Currently, it is still the best choice to predict the state of dynamic protein interaction by means of gene expression levels [15]. But, two challenges need to be tackled: (1) How to effectively integrate protein interaction data and gene expression data to predict dynamic interaction proteins? (2) How to screen out key dynamic interaction among them?

Here, we presented a dynamic PPIs’ hypothesis, that is, ***P***ermanent interactions could play a critical role in maintaining physiological processes of normal cells, such as immune system, once it changes dramatically in***to T***ransient interactions from normal conditions to pathologic conditions (**P2T**) and long-lasting interaction is blocked, it results in cellular dysfunction or diseases. Correspondingly, ***T***ransient interactions could be involved in signal transduction, like cell proliferation, a sustaining proliferative signaling leads ultimately to uncontrolled growth of the cells when it changes in***to P***ermanent interactions from normal conditions to pathologic conditions (**T2P**). Based on the dynamic PPIs’ hypothesis, we first applied a new approach, PPI-express, to predict key genes for 18 cancers. Then we defined the transient and permanent interactions under normal and cancer conditions, and further detected the dynamic interaction pairs that are dynamically switching between the two conditions. Finally, we mapped the key cancer genes (KCG) into dynamic interaction pairs to filter out the key dynamic interactions (KDI) of cancer. Two types of KDI pairs were analyzed, and our findings could help to explain a few hallmarks of cancer. About 450 human microarray datasets comprising of more than 40,000 samples (arrays) were involved in this study.

## Results

### Application on 18 cancer datasets of PPI-express

The results of the method evaluation suggested that PPI-express is a promising approach to identify cancer-associated genes comparing with NGP and GraphRank (see Supplementary file 1 for the performance comparison of three methods). To detect reliable KCG for further screening of KDI pairs (Figure 1 shows its flowchart), PPI-express was applied to the datasets of 18 different cancers (Supplementary Table S1). Based on them and the human PPI data, 18 ranked gene lists were identified and top 5% of them were defined as KCG. It contained about 140 genes in each cancer dataset and a total of 1,474 genes were involved. 93 genes were found at least 5 times in all types of cancer, and two cell cycle-related genes, CDK1 and CCNA2 (Cyclin A2), appeared 13 times which was the highest frequency of occurrence (lower left in Figure 2A). They frequently involve tumor-associated mutations [16], and the cyclin B/CDK1 and the cyclin A/CDK2, two specific subset of CDK-cyclin complexes, directly control cell cycle progression in G2 and G1 phases [17,18]. We also found that the intersections of the gene sets were significantly larger than random expectations, and the higher their frequencies, the greater the significant of intersection by 1000 permutations (upper right in Figure 2A). Supplementary Table S2 lists more details of KCG.

**Fig 1.**
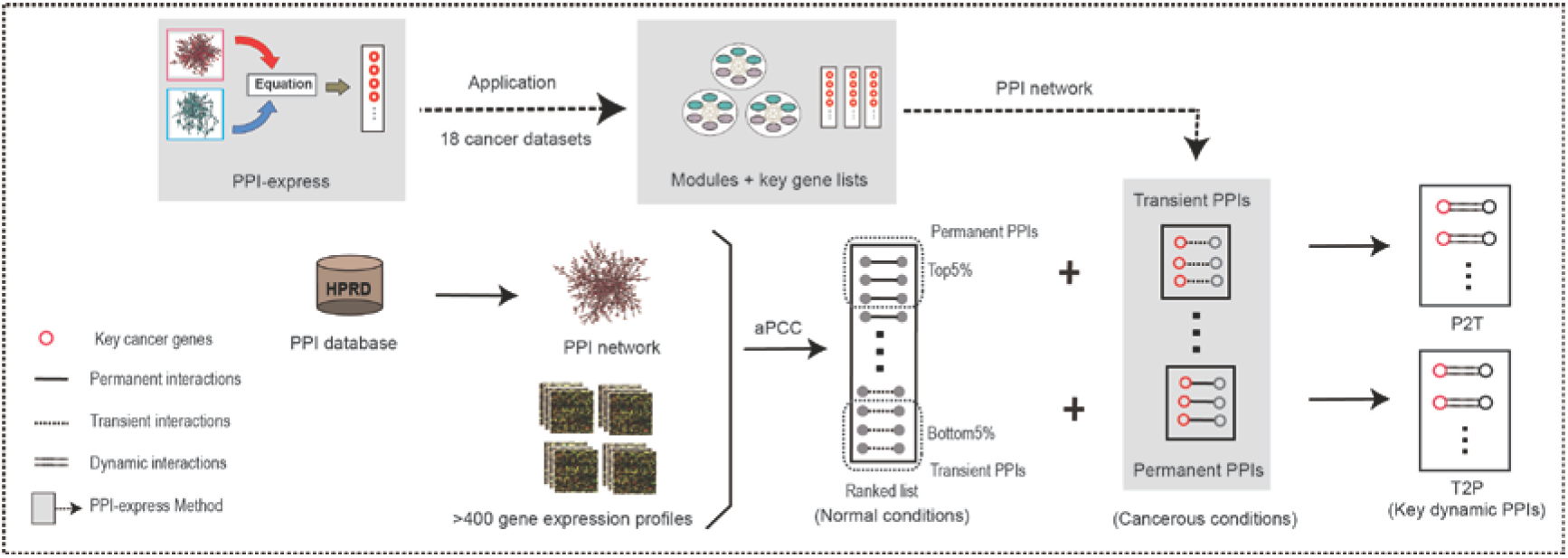
The wokflow of identification of KDI. aPCC: average Pearson Correlation Coefficient.

**Fig 2.**
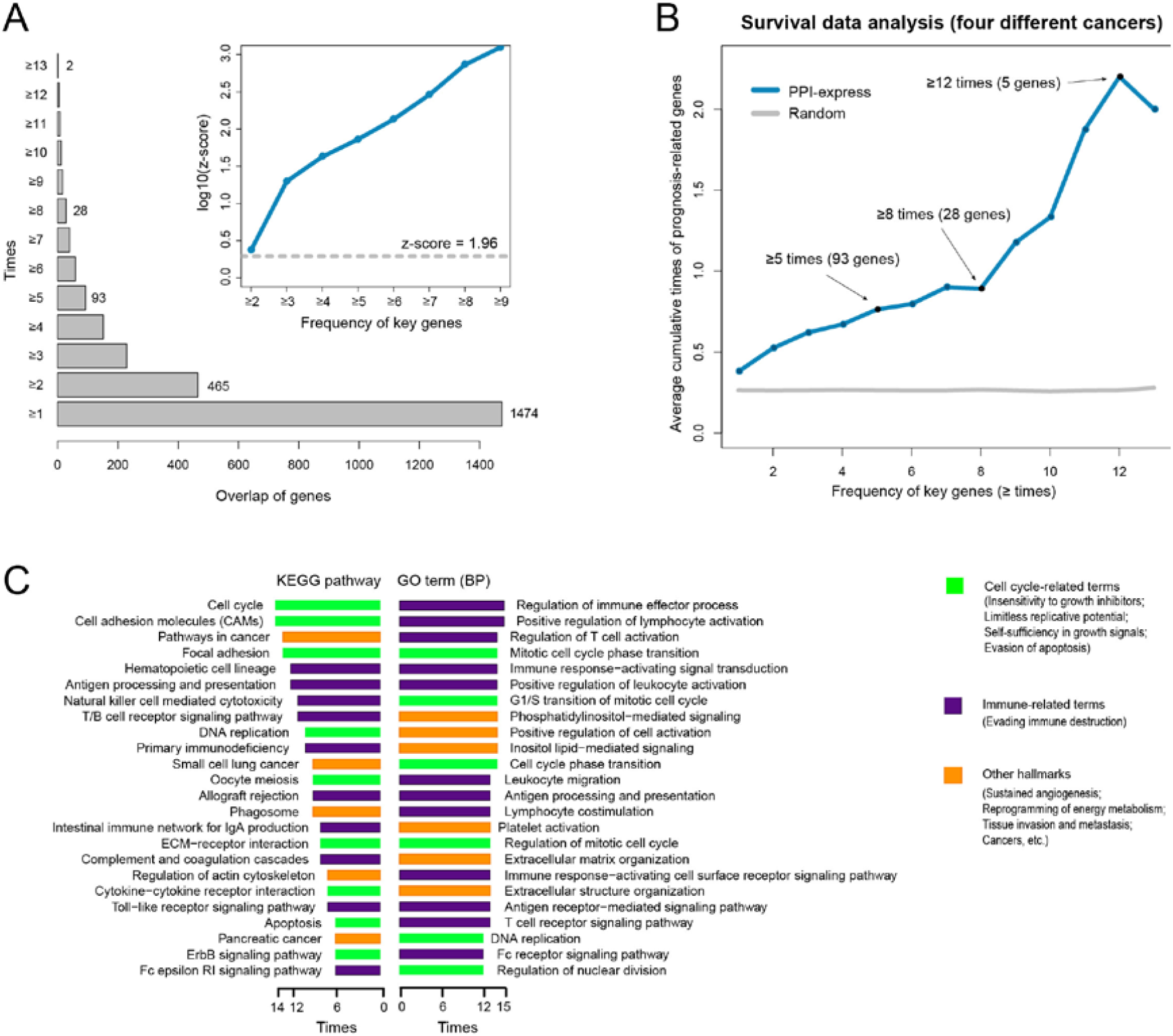
Analysis of key cancer genes. (A) Overlaps of predicted KCG from 18 cancer datasets(lower left). We calculated the z score by a thousand times simulation for estimating the intersections of gene sets with same sizes (there is no common gene when the number of overlap simulation is more than 9 gene sets), and horizontal dashed line at z score value of 1.96 indicates a significant p value in the upper-right region. (B) The average cumulative times of prognosis-related genes in all KCG with different frequency (i.e. times ⩾1, 2 … 13) based on 4 survival datasets. The random (averaging values of sampling randomly for 1000 times) is calculated as a reference. It implies that the higher frequency of KCG, the tighter correlation between them and cancer. (C) The overall results of KEGG pathway and GO enrichment analysis based on KCG by Fisher test. The terms with higher frequency were listed from 18 cancer datasets (KEGG ⩾6 times and GO (BP) ⩾12 times). They may be artificially divided into three categories.

To confirm that KCG is closely related to cancer outcomes, we counted times which each prognosis-related gene repeatedly appears in 4 independent datasets with survival data. As shown in Figure 2B, we found that, compared to the random, KCG contained more prognostic genes, and the times gradually increase with the gradual increase of KCG frequency, hinting that KCG is indeed closely related to cancer. In addition, we also manually searched “high-frequency” KCG using PubMed searches and found that most of them were often described as several striking keywords, such as a potential biomarker and a therapeutic target, in the article’s title or abstract, like, CDC20 [19] and BRCA1 [20].

To investigate the function of KCG, we performed functional enrichment analysis for 18 sets of KCG based on GO (BP) terms and KEGG pathways database by using Fisher test. The results consistently showed that KCGs were closely linked to Cell cycle, Immunodeficiency, Apoptosis and Angiogenesis which were reported in the literature as hallmarks of cancer [21], suggesting that they are involved in many cancer-related biological pathways. We then counted the number of pathways, which are consistently enriched in KCG of 18 cancers, and found that the high frequency pathways were focused on the cell cycle-related and immune-related pathways (Figure 2C). The former includes “cell cycle”, “DNA replication”, “cell division”, etc., while the latter contains “primary immunodeficiency”, “B/T cell receptor signaling pathway”, etc. Evidently, these functions were inextricably linked with cancer[22,23]. Other hallmarks, such as Angiogenesis, Pyrimidine/Purine metabolism, and Regulation of protein kinase B signaling cascade, were also significantly enriched in KCGs across multiple cancers (Figure 2C).

### Identification and analysis of KDI

#### Definitions and network properties of permanent and transient PPI pairs

We further predicted KDI based on KCG derived from the application of PPI-express to multiple cancers. The workflow of predicting KDI was shown in Figure 1. Besides KCG, the following two parts were indispensable for it: (1) Transient and permanent interactions under normal conditions were screened out from a large number of non-cancer gene expression datasets; (2) Two types of interactions under cancer conditions were obtained based on the significant up-regulation or down-regulation modules. The dynamic switching of protein interactions from normal to cancer conditions includes P2T and T2P PPI pairs (Figure 1).

There were a total of 36,746 PPI pairs in the HPRD database and 36,124 PPI pairs coexisting in more than 100 datasets of the 412 non-cancer datasets. Using average PCC (aPCC) of gene pairs to rank PPI pairs in these datasets, we confirmed 1,806 pairs for permanent and transient PPI pairs under normal conditions, respectively. GSTM1 (glutathione s-transferase M1) and GSTM2 (glutathione s-transferase M2) pair ranked highest (aPCC: 0.756) in the permanent PPI pairs, while ARHGDIA (Rho GDP-dissociation inhibitor 1) and CDKN1B (Cyclin-dependent kinase inhibitor 1B) pair ranked lowest (aPCC: -0.267) in the transient PPI pairs, which means a negative correlation under normal conditions with no interaction. Permanent and transient PPI pairs showed different properties in the protein interaction network. Edge betweenness (EB) is an important indicator for edges in the network reflecting the degree involved in signal transduction [24,25]. As shown in the left of Figure 3A, EB of transient PPI pairs are significantly higher than those of permanent PPI pairs, while EB of background lie in between, with statistically significant difference to both of them. The difference of two PPI pairs was consistent with previous report [15].

**Fig 3.**
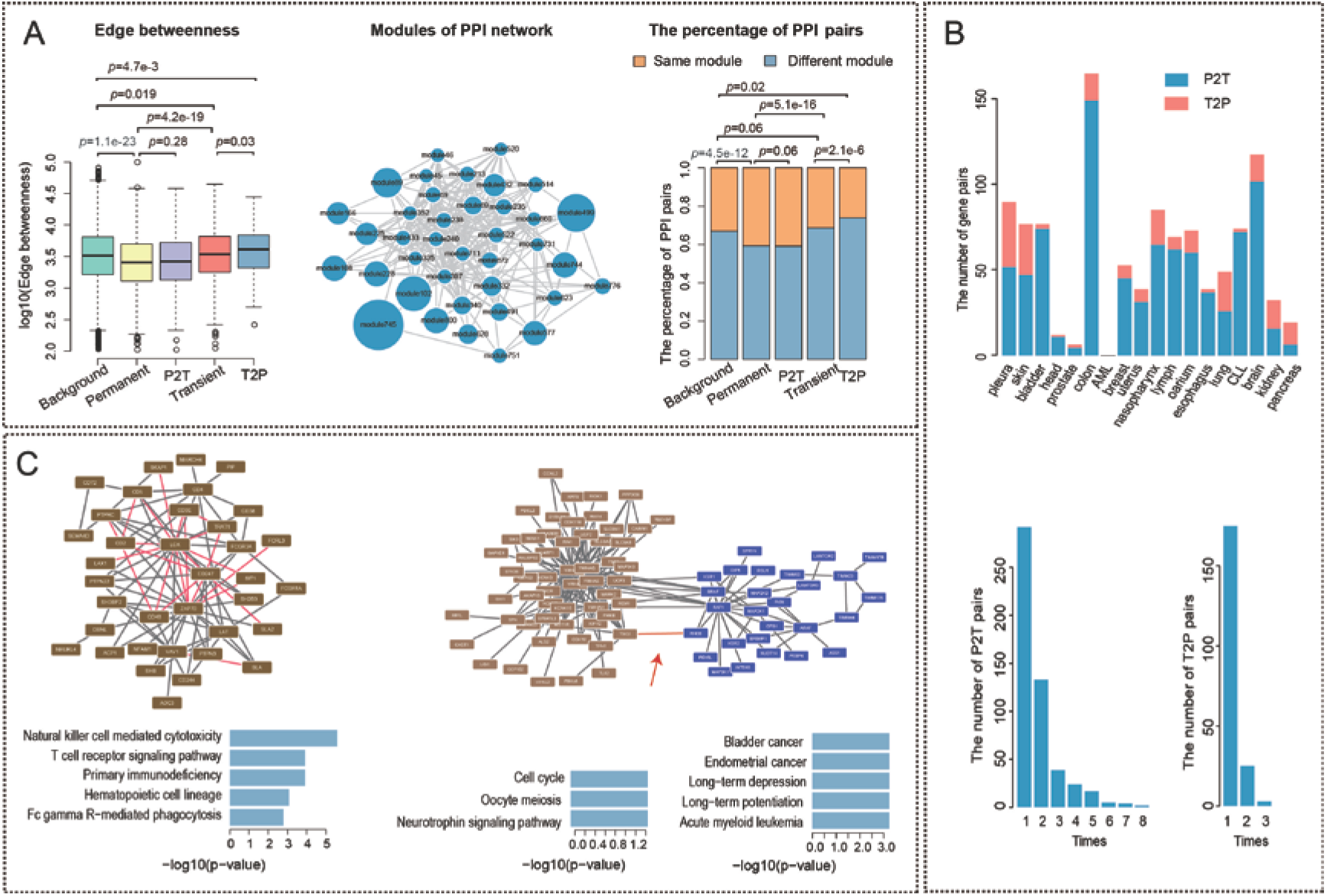
Identification and analysis of KDI. (A) Network characteristics of different PPIs (Permanent, P2T, Transient, T2P, and Background). Its left part is edge betweenness of different PPIs (p values based on one-sided Wilcoxon rank sum test), its right part is the distribution of PPIs within or between modules (p-value generated from the one-sided prop.test), and HPRD network is divided into modules with different size by iNP algorithm in the middle (The size represents the number of genes within modules, the modules containing more than 30 genes are only showed). (B) The number of T2P and P2T for 18 cancer datasets (in the upper part), and the frequency of P2T and T2P in the 18 datasets (in the lower part). (C) Function enrichment analysis of modules containing dynamic PPIs. The module including the largest number of P2T is analyzed in its left part, and two modules which is involved with T2P and can be both enriched in certain function are showed in the right part. The red lines represents the interactions between the dynamic protein pairs in the module or between modules. Enrichment analysis is performed based on KEGG pathway database by one-sided Fisher’s exact test, and top5 functions are listed.

To validate that permanent PPI pairs tend to exist intra-module, while transient PPI pairs tend to exist inter-modules, for higher EB contributed to communication of different funtional modules the phenomenon, we partitioned the HPRD network into 951 modules using iNP algorithm [26], with 39 modules containing more than 30 genes (the middle of Figure 3A). As shown in the right of Figure 3A, the proportion of PPI pairs between modules in transient PPI pairs is significantly higher than that in permanent PPI pairs and background; and this proportion in permanent PPI pairs is significantly lower than that in background (using single-tail proportion test). Therefore, we inferred that compared with permanent PPI pairs, transient PPI pairs tend to play signal transduction roles between modules.

We next investigated EB of KDI pairs. The difference of EB between T2P KDI pairs and P2T KDI pairs is larger than transient and permanent PPI pairs. And T2P KDI pairs had higher EB than transient PPI pairs, while we didn’t detect a significant difference between permanent PPI pairs and P2T KDI pairs (the left of Figure 3A); When putting T2P and P2T KDI pairs into the partitioned functional modules of the HPRD network, we found that just like transient PPI pairs, T2P KDI pairs also tended to locate between modules, and the percentage of pairs belonging to different modules was significantly higher than that in transient PPI pairs (p<2.1e-6); while the difference was not obvious between P2T KDI pairs and permanent PPI pairs (the right Figure 3A). Results indicated that T2P KDI pairs may play a more pivotal role in connecting different functional modules.

We found that KDI was heterogeneous. Firstly, we observed that the number of KDI pairs in 18 cancer datasets was from 0 to 127 (no KDI pair was found in the Acute Myelocytic Leukemia dataset). The number of average P2T KDI pairs was 51 and 13 for T2P KDI pairs, and the distributions of KDI pairs in different cancer types were shown in the upper panel of Figure 3B. After combination, we got 516 P2T KDI pairs and 203 T2P KDI pairs. P2T KDI pairs outnumbered T2P KDI pairs for most cancers, probably because many functions of normal cells need to be silenced in cancer cells, but a few functions such as cell proliferation need to be reinforced. Secondly, we examined the frequency of two types of KDI pairs in different cancers. As shown in the bottom panel of Figure 3B, the frequency of P2T KDI pairs is much higher than T2P KDI pairs. In P2T pairs, two pairs appeared in 8 cancer datasets as the highest frequency, one of them was HBA2 and HBB pair (two subunits of Hemoglobin). The dramatic dynamic change of HBA2 and HBB in blood would influence the function of this heterodimer (oxygen transport), which can lead to hypoxia, activating a series of hypoxia regulating proteins (such as galectin-1) and influenced cancer process and patient prognosis [27]. Besides, Woong-Shick et al [28] found that the content of Hb-α (HBA) and Hb-β (HBB) were significantly different in ovarian cancer serum samples by an experimental method, and implied that they could be used as diagnostic biomarkers. Based on the above evidences, the dynamic change of HBA2 and HBB may be related with the cancer process. In T2P pairs, HDAC2 (Histone deacetylase 2) and STAT3 (Signal transducer and activator of transcription 3) appeared in 3 cancer datasets as the highest frequency.

#### Different pathways affected by P2T and T2P

We subsequently investigate KDI’s influence on the biological processes of cancer cells. As for P2T, we mainly focused on the enriched functions of modules containing most P2T pairs. While for T2P we focused on the respective enriched functions of two modules connected by T2P pairs. The left of Figure 3C listed the enriched functions of the module containing most P2T. There are 36 nodes including proteins from 23 P2T in it, and the most enriched functions are natural killer cell mediated cytotoxicity and T cell receptor signaling pathway, which have something to do with the body immunological functions, such as LCK and ZAP70 [29]. The right of Figure 3C showed two modules connected by a T2P KDI pair, RHEB and TSC2, and their respective functions. We chose these two modules because they both can enrich significant functions. The most enriched significant functions were cell cycles and some cancer pathways (the right of Figure 3C). Because modules containing P2T maintained the normal biological functions, the silence of P2T KDI pairs’ interactions may have a key effect in silencing the biological functions. As for T2P, because there was no expression of their genes in normal cells, there was no communication between the modules connected by T2P, as well. However, in cancer cells, the activation of their interaction states would lead to the continuous information passing between the modules connected by T2P, making uncontrolled growth of cancer cells as a result. Identification of KDI was meaningful for us to design relevant drugs to prevent protein interaction so as to prevent the growth signal of cancer cells.

Above all, we hereby inferred that, P2T KDI pairs tending to locate within functional modules may play roles in maintaining the normal biological functions in cells, such as cell immune functions, while T2P KDI pairs tending to locate between functional modules may play roles in biological signal transduction [12], such as cell growth. In cancer cells, P2T KDI pairs tended to transfer the interaction ways from permanent interaction to transient interaction, while T2P KDI pairs reflected the continuous activation of infrequent signal transduction pathways such as the uncontrolled growth of cancer cells. These inferences were consistent with our dynamic PPIs’ hypothesis.

### Case study of P2T and T2P

We selected two cases of KDI to clearly demonstrate the dynamic PPIs’ hypothesis in cancer. The first case as a typical example of P2T KDI pairs with high frequency was FOS and JUN, which presented 7 times in 18 cancer datasets. Together with the other 11 P2T pairs in a module containing 32 genes, the most enriched function of the module was MAPK signaling pathway (Figure 4A). FOS and JUN coexist as two major components of activating protein 1 (AP-1 complex) in normal situation. As a transcription factor of heterodimer, AP-1 combines specifically with DNA and assist to activate the expression of genes related with immunity. Studies have shown that many other factors such as protein interaction can change the activity of the compound [30,31]. Among them, JUN, FOS as well as NF-κB are essential for the transcription of immune response genes, e.g. IL-2 gene[32]. In our results, JUN and FOS often participate in B/T cell receptor signaling pathways, which has been mentioned in the enriched functions of the module containing most P2T(Figure 4A). This showed that the blockage of this immune related signal pathway may affect the progress of cancer.

**Fig 4.**
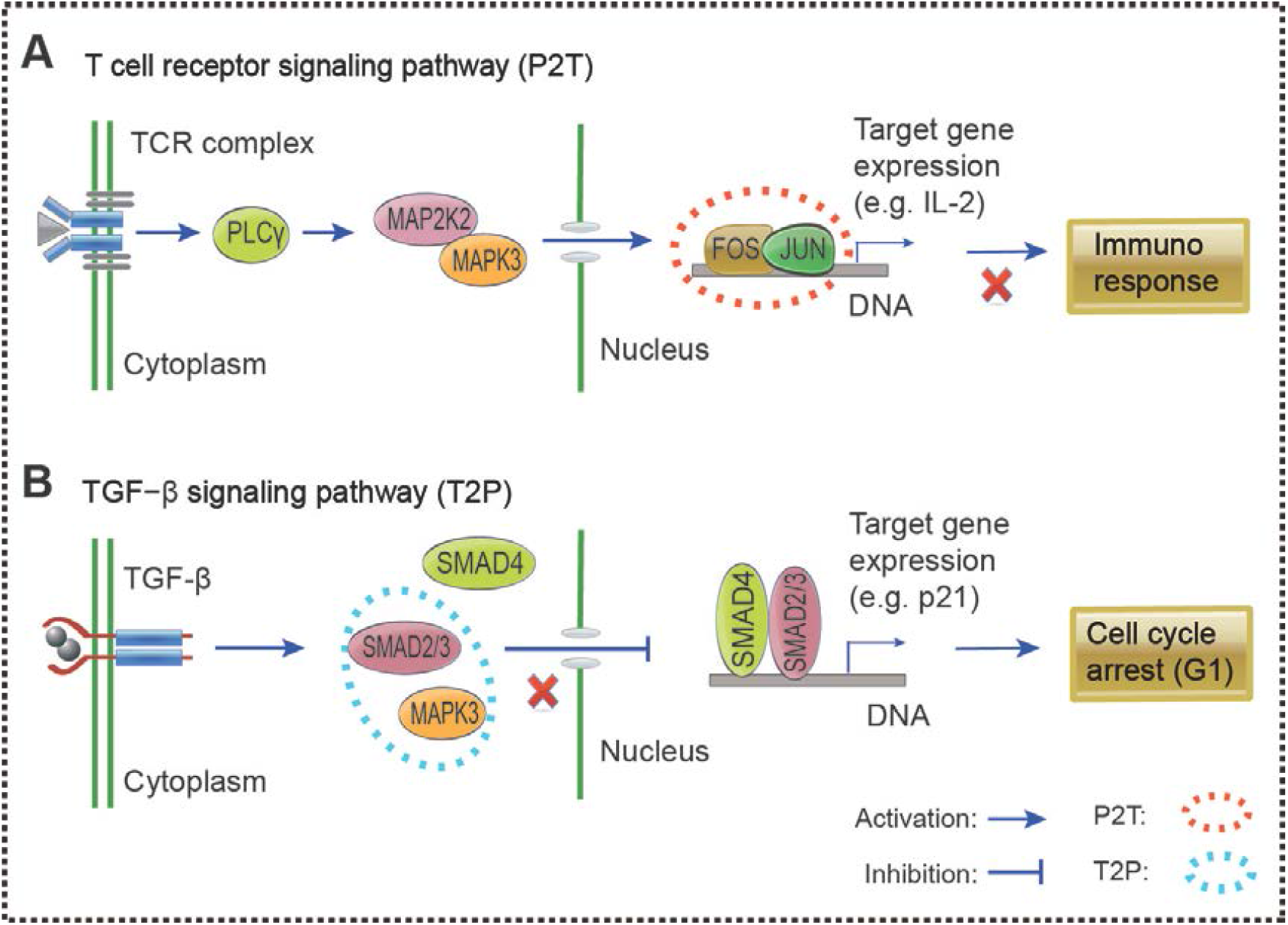
Two cases of P2T and T2P for demonstrating their mechanism based on T cell receptor signaling pathway (A) and TGF−β signaling pathway (B).

For a case in T2P, we selected SMAD2/3 (SMAD2 and SMAD3) and MAPK3. SMAD2/3 acts in a canonical SMAD signal pathway involving in cell proliferation. Through the protein interaction of SMAD family, transforming growth factor-β (TGF-β) regulates cell growth, differentiation, and apoptosis. SMAD2/3 is activated by TGF-β receptor kinases, and forms heterotrimeric complex with SMAD4 (in fact, SMAD2 and SMAD4 are two tumor suppressor genes), they then migrate into the nucleus and regulate the transcription of target genes. During this period, SMAD2/3 interacts with TbRI and SARA for its phosphorylation and complex formation, and it is subsequently separated before the activation of target genes [33]. It is called the canonical SMAD-signaling pathway which is a representative example of signal transduction [2]. A cell-cycle inhibitor, p21, can be induced by SMAD signals as a critical regulator of G1-phase progression so that cell cycle arrest can be induced [26]. As shown in Figure 4B, SMAD2/3 and MAPK3 form a T2P KDI pair based on our predictions, thereby preventing the formation of SMAD2/3 and SMAD4, and results in cell cycle progression if p21 is not induced. Other report has showed that MAPK3 belongs to ERK signaling which is an important part of MAPK signaling pathway involved in several fundamental cellular processes [34]. Interestingly, it has been reported that ERK may prevent anti-proliferative effects of TGF-β by the disruption of SMAD accumulation in the nucleus [35]. In short, the dynamic binding of SMAD2/3 and MAPK3 in cancer motivated the blockage of normal cell cycle arrest, and further leaded to uncontrolled cell growth.

### Different network structure of three types of genes in KDI

We also investigated network properties and possible influenced functions of genes forming those pairs. We separated these genes into three gene sets: only P2T gene set, only T2P gene set and common gene set. The gene numbers of only P2T gene set, only T2P gene set and common gene set were 521, 221 and 80, respectively (the upper right corner of Figure 5A). By examining their degree and betweenness, we found significant differences among them (all p-values< 0.05, KS test). That is, common gene set, only T2P gene set and only P2T gene set in the order of their most degree (Figure 5A), and the same order for betweenness (Figure 5B).

**Fig 5.**
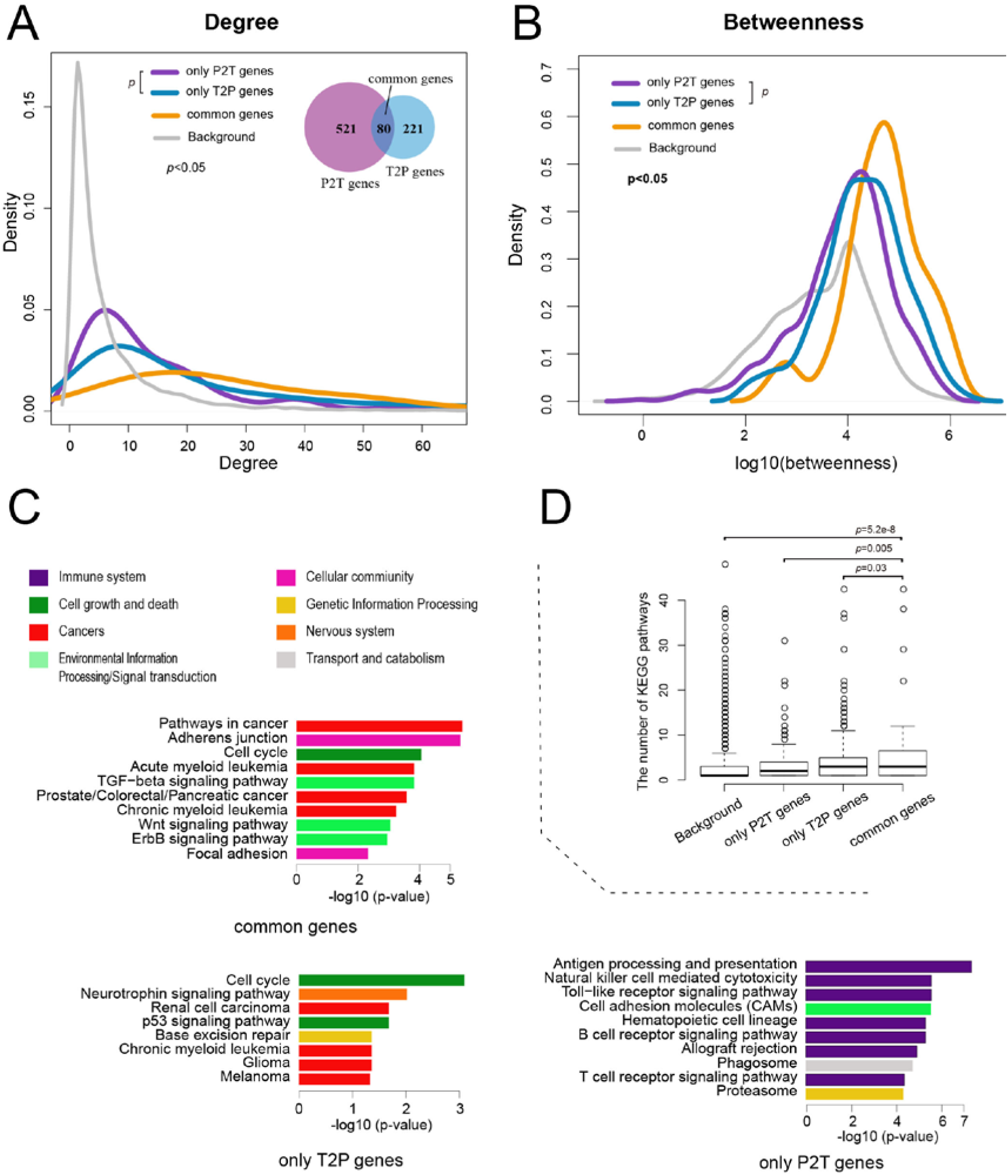
The characteristics of three gene sets involved in dynamic protein pairs. Degree(A) and Betweenness(B) of network, their function(C), and the number of KEGG pathway with their participation(D) were shown, and the venn graph for overlap of their sets was located at the upper right of A. PPIs or gene sets of HPRD were used as background and the p-values were estimated using the KS test in A, B and D. In C, KEGG-based pathway enrichment analysis of three gene sets were estimated using one-sided Fisher’s exact test and FDR-adjusted p-value. Top 10 terms of each result are shown (only T2P genes have only eight terms), and different colors represent different major types of pathways based on KEGG database.

We further performed KEGG pathway enrichment analysis for functions of three gene sets (Figure 5C). Intriguingly, we found that only P2T gene set enriched with immune-related pathways, such as antigen processing and presentation; only T2P gene set was mainly enriched for the cell cycle and neurotrophin signaling pathways, while various signal transduction pathways and other pathways such as Wnt signaling pathway, TGF-β signaling pathway and endocrine system pathways were involved in common gene set. In short, the enriched biological pathways of three types of gene sets presented a certain of preference, and the enriched functions were closely related with cancer [36]. Among them, compared with the other types of gene sets, common gene set may play more roles in completing all kinds of biological processes, which was consistent with the investigation results of network properties. In addition, we counted the number of KEGG pathways that three types of gene sets involved in (Figure 5D). We found that, compared with the other two types, common gene set involved in the most KEGG pathways, such as MAPK3 with 77 different KEGG pathways. MAPK3 plays a pivotal role in MAPK signaling pathway [35].

Compared with genes in P2T KDI pairs, genes in T2P KDI pairs could be better suited for the cancer therapy target, because the situation that transient PPI pairs transfer to permanent PPI pairs may be reversed more likely by new competitive molecule inhibitors. For example, two KDI pairs encoded by HDAC2 and HMGB1 showed a high frequency in T2P KDI pairs (7 and 8 times, respectively). The competitive inhibitors against them will directly affect the permanent PPI pair formation with other proteins. Studies have shown that PXD101/Belinostat (competitive inhibitor of HDAC2) [37] and Glycyrrhizin (competitive inhibitor of HMGB1) [38] can inhibit the cancer cell proliferation and metastasis.

## Discussion

For functional analysis, we found that two broad categories of biological function, “cell cycle-related” and “immune-related” functions were clearly dominant characteristics of KCGs rather than other cancer hallmarks in multiple-cancer application of PPI-express. Subsequently, the functions of only T2P and only P2T gene sets were also focused on cell cycle and immune. For network properties of KDI pairs, we found that permanent PPI and P2T KDI pairs tended to locate within the modules, while transient PPI and T2P KDI pairs tended to connect modules. Whether they were functional or network characteristics, these findings were consistent with our dynamic PPIs’ hypothesis.

The prediction of KDI in cancer was equivalent to placing the predicted KCG in a higher dimension, not only to the genes themselves, but also to other genes that interact with them so that we can better understand their mechanism of action. Further, a question may be asked, what promotes the formation of dynamic PPI pairs? Just like dissociation and association of general complex, dynamic switching may be attributed to two potential factors, physiological conditions and environment [6]. For example, physicochemical features of their interfaces belonged to the former, and genetic mutations can lead to the changes of their features (e.g., tumor suppressor gene, SMAD4). For the latter, competitive inhibitor can block the formation of target protein complex by binding to one of them [3]. Actually, they often occur in cancer cells [39].

In fact, the ultimate goals of cancer research are treating cancer by various treatments. Similarly, we expect our findings can provide help for understanding and explaining a few hallmarks of cancer. Specifically, two dominant functional characteristics of KCG demonstrate the general characteristics of cancer. And it may provide some enlightenment for us on cancer therapeutics, such as immunotherapies or chemotherapies. Secondly, we first studied the dynamic interaction of cancer on a large scale, dynamic PPIs’ hypothesis presented a possible cancer pathogenesis from the perspective of protein-protein interaction, and we also found KDI pairs’ gene sets with high frequencies possess higher proportions of approved drug targets. It could make cancer treatment more targeted and effective to prevent the occurrences of P2T or T2P.

In summary, we applied PPI-express proposed to 18 cancer datasets after confirming that it was superior to other competing methods. Then, KCGs were used to further discover KDI pairs, and they provided us with a possible explanation for some of cancer hallmarks. However, it also existed shortcomings, for example, we can only use the changes of gene expression to represent the dynamic switching in their corresponding proteins for the lack of the data of protein dynamic interaction, and it was the lack of consideration of different stages of cancer in the application, and definitions of transient and permanent PPIs could be too strict and uniform, etc. Nevertheless, the results and findings also have important implications for understanding and treatment of cancer in this study.

## Methods

### PPI-express and its applications

#### PPI-express, its evaluation and application

To obtain key cancer genes, we proposed a new method, ***PPI-express***, which was an improved and corresponding version Met-express [40] to prioritize cancer-associated genes by the integration of ***PPI*** network and gene co-***express***ion network. Supplementary file 1 was provided for more details of *PPI-express* and method evaluation.

We collected18 cancer datasets including 659 cancer samples and 513 normal samples for PPI-express’s application only from DataSets (GDS) of GEO based on the rigorous criteria. (Supplementary Table S1).

#### Analysis and validation

For functional enrichment analysis, no inferred from electronic annotation (no IEA) biological process of gene ontology (GO, http://geneontology.org/) and KEGG pathways (http://www.genome.jp/kegg/) were enriched in each KCG lists of 18 cancer datasets using Fisher’s exect test (fdr-adjusted p < 0.05).

As for KCG lists of 18 datasets, we concluded that the more times KCG presented in cancer datasets, the more related with cancer. Validation of this trend will indirectly verify our predictions. To validate it, we examined that high-frequency KCG contains the proportion of prognosis genes (two significant p-values, see Supplementary file 1 for more details) based on survival analysis of datasets of four types of cancers.

### Detection and analysis of KDI

#### Identification of transient and permanent PPIs under normal conditions

we screened human non-cancer microarray data with 10 to 1,000 sample size and eliminated sources from stem cells and cancer in GEO DataSets using its advanced search tool. A total of 412 human non-cancer datasets containing gene numbers larger than 3,000 were obtained after data normalization.

For each PPI pair in HPRD, gene pairs encoding the PPI pairs coexist in more than 100 datasets among 412 non-cancer datasets were screened out. PCCs of their expression values of these screened pairs were calculated in each dataset and average PCCs (aPCCs) of each gene pair in all datasets were obtained. All of PPI pairs were then ranked by their aPCCs and top 5% were considered as permanent PPI pairs, also with bottom 5% as transient PPI pairs under normal conditions.

#### KDI prediction

The dynamic interaction of cancer proteins assumed that the types of protein interaction are reversible in cancer. By means of intermediate step of the preceding KCG prediction, AUC (area under ROC curve)of gene co-expression modules contained gene pairs were used for the prediction of dynamic interaction pairs. Here, we only focused on significantly up-regulated modules with AUC greater than 0.8 and down-regulated modules with AUC less than 0.2. We proposed a T2P KDI pair as a transient PPI pair appearing in a significantly up-regulated module, which represented a transient state from low expression (no interaction) under normal conditions to high expression (likely interaction) in cancer. Vise versa, a P2T KDI pair was denoted as a permanent PPI appearing in a significantly down-regulated module, which represented a permanent state from high expression (close interaction) under normal conditions to low expression (no interaction) under cancer conditions. After prediction of T2P and P2T dynamic interaction pairs, we defined a key cancer dynamic interaction pair (KDI) of permanent/transient PPI pairs if at least one member in the dynamic interaction pair belonged to KCGs (Figure 1).

#### Analysis of KDI

For network topology, edge-betweenness (EB) of KDI pairs, degree and betweenness of genes in the KDI pairs were examined respectively[41]. R package igraph was used for calculation of three topological features and iNP, an iteration algorithm for network partition with high accuracy and function consistency[26], was used for network partition. Cytoscape v3.0.1 software (http://www.cytoscape.org/index.php) was used for network display. We selected HPRD as the background network when performing KEGG pathway enrichment.

### Other Implementation Details

To standardize gene names across microarray platforms, we mapped the gene names to the official gene symbols by a mapping table derived from HGNC database [42]. All datasets above were downloaded in October, 2018, and their lists and specific selection criteria can be found in the Supplementary Table S1 and file 1.

## Acknowledgements

This work was supported by National Natural Science Foundation (31501461); Anhui Science and Technology Major Project (no.18030701189); Huainan science and technology project (2017A0421); the Key Support Program for Outstanding Young Talents in University of Anhui Province (no. gxyqZD2016264).

## Supporting information

S1 File. PPI-express and method evaluation; S1-1 Table. The information of 18 cancer datasets and their selection criteria; S1-2 Table. The list and three selection criteria of 412 non-cancer datasets; S2-1 Table. The number of key cancer genes of each dataset in this study; S2-2 Table. All key cancer genes and their frequencies which appear in 18 sets of key cancer gene.

## References

1. Masoudi-Nejad A, Meshkin A, Haji-Eghrari B, Bidkhori G (2012) Candidate gene prioritization. Molecular genetics and genomics : MGG 287: 679–698.

2. Barrios-Rodiles M, Brown KR, Ozdamar B, Bose R, Liu Z, et al. (2005) High-throughput mapping of a dynamic signaling network in mammalian cells. Science 307: 1621–1625.

3. Ozbabacan SE, Engin HB, Gursoy A, Keskin O (2011) Transient protein-protein interactions. Protein engineering, design & selection : PEDS 24: 635–648.

4. Nooren IM, Thornton JM (2003) Diversity of protein-protein interactions. The EMBO journal 22: 3486–3492.

5. Scheffzek K, Ahmadian MR, Kabsch W, Wiesmuller L, Lautwein A, et al. (1997) The Ras-RasGAP complex: structural basis for GTPase activation and its loss in oncogenic Ras mutants. Science 277: 333–338.

6. Nooren IM, Thornton JM (2003) Structural characterisation and functional significance of transient protein-protein interactions. Journal of molecular biology 325: 991–1018.

7. Przytycka TM, Singh M, Slonim DK (2010) Toward the dynamic interactome: it’s about time. Briefings in bioinformatics 11: 15–29.

8. Chen B, Fan W, Liu J, Wu FX (2014) Identifying protein complexes and functional modules--from static PPI networks to dynamic PPI networks. Briefings in bioinformatics 15: 177–194.

9. Mishra GR, Suresh M, Kumaran K, Kannabiran N, Suresh S, et al. (2006) Human protein reference database--2006 update. Nucleic acids research 34: D411–414.

10. Szklarczyk D, Morris JH, Cook H, Kuhn M, Wyder S, et al. (2016) The STRING database in 2017: quality-controlled protein–protein association networks, made broadly accessible. Nucleic acids research: gkw937.

11. Ruan J, Zhang W (2008) Identifying network communities with a high resolution. Physical review E, Statistical, nonlinear, and soft matter physics 77: 016104.

12. Taylor IW, Linding R, Warde-Farley D, Liu Y, Pesquita C, et al. (2009) Dynamic modularity in protein interaction networks predicts breast cancer outcome. Nat Biotechnol 27: 199–204.

13. Liu Y, Koyuturk M, Barnholtz-Sloan JS, Chance MR (2012) Gene interaction enrichment and network analysis to identify dysregulated pathways and their interactions in complex diseases. BMC systems biology 6: 65.

14. La D, Kong M, Hoffman W, Choi YI, Kihara D (2013) Predicting permanent and transient protein-protein interfaces. Proteins 81: 805–818.

15. Das J, Mohammed J, Yu H (2012) Genome-scale analysis of interaction dynamics reveals organization of biological networks. Bioinformatics 28: 1873–1878.

16. Malumbres M, Barbacid M (2009) Cell cycle, CDKs and cancer: a changing paradigm. Nature reviews Cancer 9: 153–166.

17. Kastan MB, Bartek J (2004) Cell-cycle checkpoints and cancer. Nature 432: 316–323.

18. Sherr CJ (1996) Cancer cell cycles. Science 274: 1672–1677.

19. Kidokoro T, Tanikawa C, Furukawa Y, Katagiri T, Nakamura Y, et al. (2008) CDC20, a potential cancer therapeutic target, is negatively regulated by p53. Oncogene 27: 1562–1571.

20. James CR, Quinn JE, Mullan PB, Johnston PG, Harkin DP (2007) BRCA1, a potential predictive biomarker in the treatment of breast cancer. The oncologist 12: 142–150.

21. Hanahan D, Weinberg RA (2011) Hallmarks of cancer: the next generation. Cell 144: 646–674.

22. Vogelstein B, Papadopoulos N, Velculescu VE, Zhou S, Diaz LA, Jr., et al. (2013) Cancer genome landscapes. Science 339: 1546–1558.

23. Disis ML (2010) Immune regulation of cancer. Journal of clinical oncology : official journal of the American Society of Clinical Oncology 28: 4531–4538.

24. Radicchi F, Castellano C, Cecconi F, Loreto V, Parisi D (2004) Defining and identifying communities in networks. Proceedings of the National Academy of Sciences of the United States of America 101: 2658–2663.

25. Girvan M, Newman ME (2002) Community structure in social and biological networks. Proceedings of the National Academy of Sciences of the United States of America 99: 7821–7826.

26. Matsuura I, Denissova N, Wang G, He D, Long J, et al. (2004) Cyclin-dependent kinases regulate the antiproliferative function of Smads. Nature 430: 226–231.

27. Le QT, Shi G, Cao H, Nelson DW, Wang Y, et al. (2005) Galectin-1: a link between tumor hypoxia and tumor immune privilege. Journal of clinical oncology : official journal of the American Society of Clinical Oncology 23: 8932–8941.

28. Woong-Shick A, Sung-Pil P, Su-Mi B, Joon-Mo L, Sung-Eun N, et al. (2005) Identification of hemoglobin-alpha and -beta subunits as potential serum biomarkers for the diagnosis and prognosis of ovarian cancer. Cancer science 96: 197–201.

29. Fulop T, Jr., Gagne D, Goulet AC, Desgeorges S, Lacombe G, et al. (1999) Age-related impairment of p56lck and ZAP-70 activities in human T lymphocytes activated through the TcR/CD3 complex. Experimental gerontology 34: 197–216.

30. Eferl R, Wagner EF (2003) AP-1: a double-edged sword in tumorigenesis. Nature reviews Cancer 3: 859–868.

31. Chen L, Glover JN, Hogan PG, Rao A, Harrison SC (1998) Structure of the DNA-binding domains from NFAT, Fos and Jun bound specifically to DNA. Nature 392: 42–48.

32. Aderem A, Ulevitch RJ (2000) Toll-like receptors in the induction of the innate immune response. Nature 406: 782–787.

33. Derynck R, Zhang YE (2003) Smad-dependent and Smad-independent pathways in TGF-beta family signalling. Nature 425: 577–584.

34. Chang L, Karin M (2001) Mammalian MAP kinase signalling cascades. Nature 410: 37–40.

35. Dhillon AS, Hagan S, Rath O, Kolch W (2007) MAP kinase signalling pathways in cancer. Oncogene 26: 3279–3290.

36. Vogelstein B, Kinzler KW (2004) Cancer genes and the pathways they control. Nature medicine 10: 789–799.

37. Bantscheff M, Hopf C, Savitski MM, Dittmann A, Grandi P, et al. (2011) Chemoproteomics profiling of HDAC inhibitors reveals selective targeting of HDAC complexes. Nat Biotechnol 29: 255–265.

38. Smolarczyk R, Cichon T, Matuszczak S, Mitrus I, Lesiak M, et al. (2012) The role of Glycyrrhizin, an inhibitor of HMGB1 protein, in anticancer therapy. Arch Immunol Ther Exp (Warsz) 60: 391–399.

39. Zhang J, Yang PL, Gray NS (2009) Targeting cancer with small molecule kinase inhibitors. Nature reviews Cancer 9: 28–39.

40. Jelovac D, Armstrong DK (2011) Recent progress in the diagnosis and treatment of ovarian cancer. CA: a cancer journal for clinicians 61: 183–203.

41. Ozgur A, Vu T, Erkan G, Radev DR (2008) Identifying gene-disease associations using centrality on a literature mined gene-interaction network. Bioinformatics 24: i277–285.

42. Seal RL, Gordon SM, Lush MJ, Wright MW, Bruford EA (2011) genenames.org: the HGNC resources in 2011. Nucleic acids research 39: D514–519.

