## Supplemental file 1 for "Large-scale prediction of key dynamic interacting proteins in multiple cancers"

**Prediction of multi-cancer key dynamic interactions based on PPI-express**

**PPI-express and method evaluation**

**INTRODUCTION**

In the past decade, many computational methods for identifying disease-related genes have been developed based on different data sources [1-3]. Among them, gene expression and protein-protein interaction (PPI) data are two basic data types, and network topology is often involved in the analysis of data. Several studies were reported on the identification of disease signatures based primarily on one type of them, namely, a single PPI network data [4, 5] or co-expression network data combined or not combined with other data [6, 7]. Furthermore, based on these two types of data, many related studies have been carried out. For the combination of data, one type of data was usually used as the basic network, and it was further screened and trimmed for the discovery of candidate genes or subnetworks. An example of TF–target regulatory network was that Reverter et al. [8] developed a method called Regulatory Impact Factors (RIF) to prioritize disease genes by the change of correlation between transcription factors (TF) and their interacting differentially expressed genes. And there are many cases in which PPI network was regarded as the basic network [9, 10]. Two studies of them deserve a special mention. Wu et al. [11] developed a novel method, Networked Gene Prioritizer (NGP), to prioritize cancer-associated genes based on gene expression and PPI data. Through two screening of PPI network, NGP got subnets with high weight interactions and differentially expressed genes by calculating Spearman coefficient and T-test, respectively, and prioritized their genes using Z scores from analysis. In their study, existing methods were grouped into three main types, then, their method was compared with the Heat Kernel Ranking method[12], the method by Taylor et al.[10], and RIF [8], which were regarded as typical representatives of these types. The results showed NGP performs better than other methods by application on breast cancer and lung cancer datasets. The second method has subsequently been proposed for biomarker discovery and was called stSVM [13] whose core algorithm can be achieved by GraphRank, which is dependent on a random walk kernel and smoothing of network (graph) to process network data. Based on gene expression and PPI data, the performance of the algorithm was better than other competing methods, such as netRank [14]. However, based on gene co-expression network and PPI network, few studies have been reported on the prediction of disease genes.

The interactions between proteins (genes) may significantly alter under the cancerous conditions comparing to normal. The result leads to alterations in topological structure of certain functional subnetworks [15]. And nodes’ degrees and characteristics of modules are the highlight of the structure. So we hypothesize that the formation of new cancer-specific functional modules is more advantageous to the occurrence and development of cancer, and key cancer genes (KCG) tend to possess more co-expression partners during cancer processes (the PPI-express hypothesis). Based on this hypothesis, we propose a new approach called **PPI-express,** to prioritize cancer-associated genes by the integration of ***PPI*** Network and gene co-***express***ion Network. In fact, it is an improved and corresponding version Met-express [16], which was a method previously developed by our laboratory. In PPI-express, genes are assigned larger scores if the cancer module where they locate are more specific and they possess relatively higher degree in this module. Furthermore, the alterations of PPI may be rooted from dramatic switching between transient and permanent PPI, which are two types of protein interactions and similar features of complex interface may provide a structural basis for the switching [17].

**METHODS**

PPI-express

PPI-express was developed based on Met-express [16], which included three important steps: (1) construction of gene co-expression network and metabolic network, (2) measure of cancer-specific modules of co-expression network, and (3) calculation of the importance enzyme-coding genes. PPI-express’ procedures were generally similar to it, except the following three points: (1) Metabolic network was replaced by PPI network of human Protein Reference Database (HPRD, <http://www.hprd.org/>, Release 9) [18]. (2) For a moderately dense network, top 2 genes with the highest Pearson Correlation Coefficient (PCC) were chosen to construct the gene co-expression network rather than top 3 genes. (3) the formula of evaluating the key genes was improved. and the workflow of PPI-express was shown in Figure 1A.

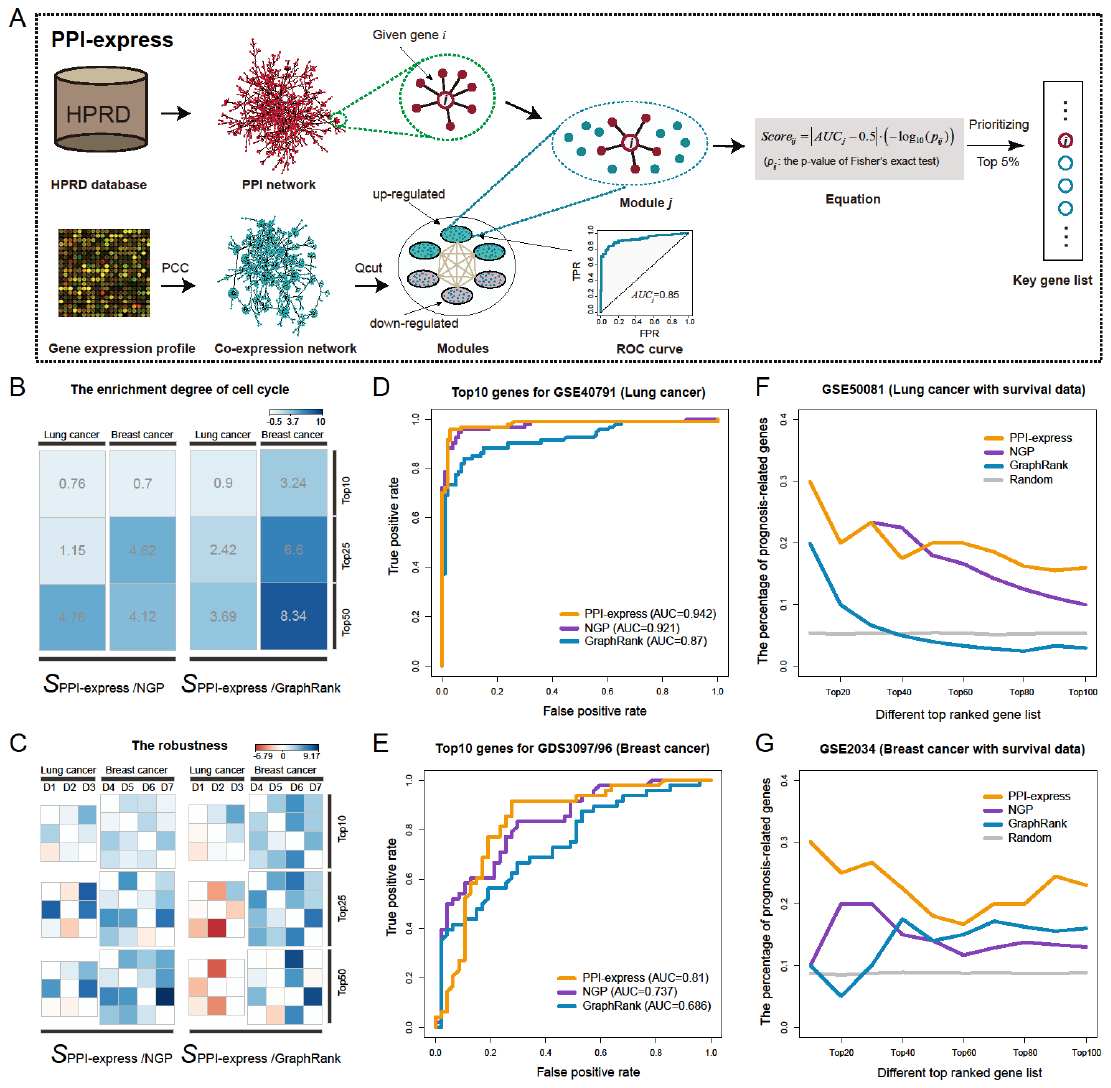

**Figure 1.** Flowchart and method comparison for PPI-express. (A) Flowchart of PPI-express. PCC: Pearson Correlation Coefficient; Qcut: a module detection algorithm. (B-G) The comparison of PPI-express, NGP, and GraphRank. (B, C) Based on top 10, 25, 50 predicted genes of each cancer, the enrichment degree of cell cycle pathway (B) and the robustness (C) were measured by KEGG pathway enrichment analysis and KS test, respectively. The results were displayed through the defined value S. That is, the smaller p value obtained by PPI-express comparing to the tested method, the more blue the appearance of the square box (the larger positive in box), conversely, the more red box (the smaller negative in box). (D, E) The classification power of genes for discriminating between normal and cancer samples. GSE40791 (D) and GDS3097/96 (E) were analyzed by SVM method based on top 10 predicted genes in common for each cancer, respectively, and AUC values (10-fold cross-validation) were displayed. (F, G) The percentage of prognosis-related genes in top predicted genes with different sizes (top 10, 20 ... 100). GSE50081 (F) and GSE2034 (G) were analyzed to assess the correlation between them and the clinical outcome comparing to the random gene sets (averaging values of sampling randomly for 1000 times). Overall, PPI-express is significantly better than other two methods.

For the formula, given a gene G_i_ in a co-expression module S_j_, the cancer-specific score of G_i_ was finally integrated as:

(1).

|*AUC_j_* -0.5| represents the degree of cancer specificity of module S*_j_*. *-log_10_(p_ij_)* measures the enrichment level of genes’ interacted partners within the module S_j_, among which *p_ij_* is the p-value from the single-tailed Fisher's exact test. Higher score of a gene means more relevant to cancer and genes of the top 5％ were chosen as the key cancer genes (KCG).

Method evaluation

For comparison of top ranked genes identified by each method, we also selected another two evaluation criteria and four corresponding datasets except for referring directly to NGP’s PPI and gene express datasets and their two criteria of method evaluation. NGP’s two criteria are the enrichment degree of cell cycle pathway of top gene sets and robustness of methods. We used Fisher's exact test for detecting functional enrichment and Kolmogorov-Smirnov (KS) test for assessing the similarity of results between different data sets of the same cancer. The other two criteria we proposed are as follows: (1) we utilized top gene sets as a classifier to discriminate between normal and cancer samples. Support Vector Machines (SVM) method was employed in Weka v3.6 (<http://www.cs.waikato.ac.nz/ml/weka/>); (2) we detected the proportion of prognosis-related genes in the top gene sets. In particular, for each single gene in top ranked genes, p-value of the log-rank test (samples are clustered into two groups), and the p-value corresponding to the coefficient of gene expression in a proportional hazard model (Cox model) were calculated. Genes with both significant p-values (p-value ＜ 0.05) was identified as prognosis-related genes. Lists of all datasets and specific selection criteria can be found in the tables below (Table 1-3).

**Table 1** The information of NGP’s datasets.

| **cancer type** | **Data id** | **the number of normal samples** | **the number of cancer samples** | **the number of all samples** |
| --- | --- | --- | --- | --- |
| lung cancer | GSE10072 | 49 | 58 | 107 |
|  | GSE19804 | 60 | 60 | 120 |
|  | GSE18842 | 45 | 46 | 91 |
| breast cancer | GSE5460 | 50 | 44 | 94 |
|  | GSE7390 | 56 | 102 | 158 |
|  | GSE21653 | 95 | 117 | 212 |
|  | GSE2034 | 77 | 209 | 286 |

**Table 2** The information of datasets for ROC curve and their selection criteria in section of method comparison

| **cancer type** | **Data id^*^** | **the number of normal samples** | **the number of cancer samples** | **the number of all samples** |
| --- | --- | --- | --- | --- |
| lung cancer | GSE40791 | 100 | 94 | 194 |
| breast cancer | GDS3097/96 | 47 | 48 | 95 |

*: Three selection criteria include 1) We give priority to the dataset mentioned elsewhere in this study if it possesses a sufficient number of samples; 2) Datasets possess cancer and normal sampleas, and possess as many as possible samples in GEO database; 3) The number of normal samples and cancer samples are equal or close.

**Table 3** The information of datasets with survival information and their selection criteria in section of method comparison.

| **cancer type** | **Data id** | **the number of samples** | **reference** |
| --- | --- | --- | --- |
| lung cancer | GSE50081 | 181 | J Thorac Oncol, Der et al, 2014 |
| breast cancer | GSE2034 | 276 | Lancet, Wang et al, 2005 |
| bladder cancer | GSE13507 | 165 | J Clin Oncol, Lee et al, 2010 |
| pancreatic cancer | GDS4336 | 45 | Clin Cancer Res, Zhang et al, 2012 |

*: Three selection criteria include 1) We give priority to the dataset mentioned elsewhere in this study if it possesses a sufficient number of samples and clear sample information; 2) Dataset possesses as many as possible samples in GEO database; 3) Dataset has been clearly reported in the literature.

**RESULTS**

The performance comparison of three methods

In the cancer-specific functional modules within a co-expression network, PPI-express is able to sort out the genes which possess a relatively significant enrichment of degree in the module compared with PPI network (regarded as a normal condition) based on the PPI-express hypothesis (the workflow shown in Figure 1A). They are considered to be closely associated with cancer in this study. To demonstrate advantages of PPI-express, we compared two other competitive methods, NGP and GraphRank, based on the data from NGP. For NGP, better results were collected according to their description (the results of breast cancer by NGP-ND model and lung cancer by NGP-NR model). To provide an impartial result, we first directly employed two criteria of NGP to assess the performance of methods. The gene sets with different sizes were analyzed to demonstrate their enrichment degree of the cell cycle pathway and robustness across independent datasets. Figure 1B shows the relative enrichment degree for NGP and GraphRank comparing to PPI-express. Correspondingly, Figure 1C illustrates the results of robustness. These results show that PPI-express overall outperforms these two methods.

Indeed, these two criterion can already measure the performance of the method to some extent. However, we can’t rely completely on it due to some potential biases. For example, the robustness estimated between two independent datasets may be more affected by the weights of invariant network data in algorithm rather than the same cancer pathogenesis. Therefore, we added two additional criteria: the classification power and the correlation with clinical outcome. For the former, top 5, 10, and 15 common genes in all independent datasets were selected to discriminate between normal and cancer samples in two new test datasets with same cancer type. For breast cancer datasets, the results show the average AUC value of PPI-express is 0.754 using a 10-fold cross-validation, and is higher than NGP and GraphRank (0.741 and 0.668, respectively). For lung cancer datasets, the average AUC value of PPI-express is 0.937, which is also the highest among three methods (for details see Table 4). Figure 1D and E show the ROC curves and AUC values of three methods using top 10 common genes. For the latter, that genes influence the clinical outcome, to a certain extent, can reflect a direct correlation between them and cancer. So prognosis-related genes were compared with top ranked genes in the same cancer type and calculated their proportions. As shown in Figure 1F and G, the proportions of gene sets of PPI-express are significantly higher than two other methods as well as the random. It implies that PPI-express is more efficiently to detect prognosis-related genes than NGP and GraphRank. Thus, we can conclude that PPI-express works significantly better than the other two, suggesting that PPI-express is a promising approach to identify cancer-associated genes.

**Table 4** All AUC values and their averages obtained from three method based on different sizes of gene sets.

| Cancer Type/Dataset ID | the number of top genes | AUC Value of PPI-express | AUC Value of NGP | AUC Value of GraphRank |
| --- | --- | --- | --- | --- |
| Lung Cancer/GSE40791 | top5 | 0.906 | 0.901 | 0.698 |
|  | top10 | 0.942 | 0.921 | 0.87 |
|  | top15 | 0.963 | 0.968 | 0.921 |
|  | mean | 0.937 | 0.930 | 0.830 |
| Breast Cancer/GDS3097/96 | top5 | 0.684 | 0.727 | 0.644 |
|  | top10 | 0.81 | 0.737 | 0.686 |
|  | top15 | 0.768 | 0.758 | 0.674 |
|  | mean | 0.754 | 0.741 | 0.668 |

REFERENCES

1.	Morrison JL, Breitling R, Higham DJ, Gilbert DR: **GeneRank: using search engine technology for the analysis of microarray experiments**. *BMC bioinformatics* 2005, **6**:233.

2.	Aerts S, Lambrechts D, Maity S, Van Loo P, Coessens B, De Smet F, Tranchevent LC, De Moor B, Marynen P, Hassan B *et al*: **Gene prioritization through genomic data fusion**. *Nat Biotechnol* 2006, **24**(5):537-544.

3.	Hutz JE, Kraja AT, McLeod HL, Province MA: **CANDID: a flexible method for prioritizing candidate genes for complex human traits**. *Genet Epidemiol* 2008, **32**(8):779-790.

4.	Xu J, Li Y: **Discovering disease-genes by topological features in human protein-protein interaction network**. *Bioinformatics* 2006, **22**(22):2800-2805.

5.	Milenkovic T, Memisevic V, Ganesan AK, Przulj N: **Systems-level cancer gene identification from protein interaction network topology applied to melanogenesis-related functional genomics data**. *Journal of the Royal Society, Interface / the Royal Society* 2010, **7**(44):423-437.

6.	Zhang J, Lu K, Xiang Y, Islam M, Kotian S, Kais Z, Lee C, Arora M, Liu HW, Parvin JD *et al*: **Weighted frequent gene co-expression network mining to identify genes involved in genome stability**. *PLoS computational biology* 2012, **8**(8):e1002656.

7.	Presson AP, Sobel EM, Papp JC, Suarez CJ, Whistler T, Rajeevan MS, Vernon SD, Horvath S: **Integrated weighted gene co-expression network analysis with an application to chronic fatigue syndrome**. *BMC systems biology* 2008, **2**:95.

8.	Reverter A, Hudson NJ, Nagaraj SH, Perez-Enciso M, Dalrymple BP: **Regulatory impact factors: unraveling the transcriptional regulation of complex traits from expression data**. *Bioinformatics* 2010, **26**(7):896-904.

9.	Chuang HY, Lee E, Liu YT, Lee D, Ideker T: **Network-based classification of breast cancer metastasis**. *Mol Syst Biol* 2007, **3**:140.

10.	Taylor IW, Linding R, Warde-Farley D, Liu Y, Pesquita C, Faria D, Bull S, Pawson T, Morris Q, Wrana JL: **Dynamic modularity in protein interaction networks predicts breast cancer outcome**. *Nat Biotechnol* 2009, **27**(2):199-204.

11.	Wu C, Zhu J, Zhang X: **Integrating gene expression and protein-protein interaction network to prioritize cancer-associated genes**. *BMC bioinformatics* 2012, **13**:182.

12.	Nitsch D, Tranchevent LC, Goncalves JP, Vogt JK, Madeira SC, Moreau Y: **PINTA: a web server for network-based gene prioritization from expression data**. *Nucleic acids research* 2011, **39**(Web Server issue):W334-338.

13.	Cun Y, Frohlich H: **Network and data integration for biomarker signature discovery via network smoothed T-statistics**. *PloS one* 2013, **8**(9):e73074.

14.	Winter C, Kristiansen G, Kersting S, Roy J, Aust D, Knosel T, Rummele P, Jahnke B, Hentrich V, Ruckert F *et al*: **Google goes cancer: improving outcome prediction for cancer patients by network-based ranking of marker genes**. *PLoS computational biology* 2012, **8**(5):e1002511.

15.	Liu M, Liberzon A, Kong SW, Lai WR, Park PJ, Kohane IS, Kasif S: **Network-based analysis of affected biological processes in type 2 diabetes models**. *PLoS genetics* 2007, **3**(6):e96.

16.	Jelovac D, Armstrong DK: **Recent progress in the diagnosis and treatment of ovarian cancer**. *CA: a cancer journal for clinicians* 2011, **61**(3):183-203.

17.	Ozbabacan SE, Engin HB, Gursoy A, Keskin O: **Transient protein-protein interactions**. *Protein engineering, design & selection : PEDS* 2011, **24**(9):635-648.

18.	Mishra GR, Suresh M, Kumaran K, Kannabiran N, Suresh S, Bala P, Shivakumar K, Anuradha N, Reddy R, Raghavan TM *et al*: **Human protein reference database--2006 update**. *Nucleic acids research* 2006, **34**(Database issue):D411-414.
